# Chromosome organization of Entamoeba histolytica and Entamoeba dispar

**DOI:** 10.64898/2026.07.06.736064

**Authors:** Tetsuro Kawano-Sugaya, Seiki Kobayashi, Akira Kawashima, Yumiko Saito-Nakano, Shinji Izumiyama, Tomoyoshi Nozaki, Kumiko Nakada-Tsukui

## Abstract

*Entamoeba histolytica* is a clinically important pathogenic eukaryote and the causative agent of amoebic dysentery. *Entamoeba dispar*, a nonpathogenic commensal species that resides in the human colon, is the closest sibling species, and serves as an appropriate comparator for genome-wide analysis. Although the genome of *E. histolytica* is approximately 26.9 Mb, and the largest known genome within the genus, that of *E. invadens*, is approximately 40.9 Mb, obtaining high-quality assemblies in this genus has remained challenging due to extensive repetitive regions, tRNA gene arrays, and aneuploidy. Here, we used PacBio HiFi sequencing to assemble the genomes of the pathogenic *E. histolytica* and the nonpathogenic *E. dispar*. We reconstructed all 36 chromosomes of *E. histolytica* and 35 chromosomes of *E. dispar*, assembling each as a single continuous DNA sequence (contig). The two species exhibited high genome-wide nucleotide similarity and conserved synteny at the amino acid level. At one end of each chromosome, we identified tRNA arrays, whereas the opposite end lacked such arrays, resulting in an asymmetric chromosomal architecture. Analysis of unique-read depth revealed widespread aneuploidy in both species: *E. histolytica* is predominantly tetraploid, whereas *E. dispar* is diploid, a conclusion further supported by SNP allele-frequency distributions. These assemblies provide a robust foundation for comparative genomics in *Entamoeba* and offer detailed insights into chromosome-end structure and ploidy.

## Introduction

*Entamoeba histolytica*, the causative agent of amoebic dysentery and liver abscess, is a clinically important pathogen that contributes to substantial morbidity and mortality in low-income and sanitation-poor countries and regions. More than fifty *Entamoeba* species have been described so far^1^, inhabiting multiple environments such as the animal intestinal tract, the human oral cavity (*E. gingivalis*)^2^, sewage (*E. moshkovskii*)^3^, and marine sediments (*E. marina*)^4,5^. Because *Entamoeba* includes morphologically similar pathogenic and nonpathogenic species, genetic diversity in the 18S rRNA gene sequence has been used to distinguish them (riboprinting)^6^. Similarly, nine distinct lineages have been identified by ribosomal sequence polymorphisms recovered from insect guts, highlighting the considerable diversity within the genus^7^.

Despite this diversity, only six *Entamoeba* genomes have been sequenced and publicly released to date: *E. histolytica*^8^, *E. dispar*^9^, *E. invadens*^10^, *E. moshkovskii*^11^, *E. nuttalli*^12^, and *E. marina*^5^. Among the eighteen assemblies currently available, fifteen were generated using Sanger methods or the 454, SOLiD, or Illumina platforms. Among *Entamoeba*, pathogenic *E. histolytica* and its nonpathogenic close relative *E. dispar* are highly prioritized. Both genomes were sequenced by Sanger methods into 1,529 scaffolds for *E. histolytica* and 12,258 scaffolds for *E. dispar*^8,9^. Although these assemblies have served as invaluable resources in *Entamoeba* genomics for decades, the extensive repetitive content and complex aneuploidy characteristics of these genomes have remained major barriers to producing high-quality reference assemblies.

Repetitive elements are abundant in *Entamoeba* genomes and often coincide with synteny breakpoints, underscoring their importance in shaping genome structure and evolution^13^. Well-characterized repeat classes in *Entamoeba* include non-LTR retrotransposons: LINEs (Long Interspersed Nuclear Elements) and SINEs (Short Interspersed Nuclear Elements). Three LINE/SINE families have been identified in both *E. histolytica* and *E. dispar*, whereas *E. invadens* possesses only a single LINE family^13^. In addition, *Entamoeba*-specific repeats, ERE1 and ERE2, have been described, with ERE2 being unique to *E. histolytica*^13^. Despite extensive profiling, the limits of sequencing technologies have long hindered comprehensive characterization of these repeat elements. We have been working to improve *Entamoeba* genome assemblies using long-read sequencing. Previous efforts include the *E. histolytica* HM-1:IMSS cl6 2001 assembly generated with PacBio RS II and Hi-C data^14^, and the *E. marina* SRT209 assembly produced with Oxford Nanopore MinION^5^. These studies substantially reduced the number of scaffolds and contigs (38 scaffolds for *E. histolytica* and 202 contigs for *E. marina*), however, the position of tRNA arrays, which are thought to reside at chromosome ends, remained unresolved.

The tRNA gene array is one of characteristics in the *Entamoeba* genomes. These arrays encode one to five distinct tRNA genes, some of which are found exclusively within arrays^15^. Certain arrays also include 5S rRNA or small nuclear RNAs (snRNAs)^15^. The tRNA arrays are generally localized to subtelomeric or telomeric chromosome regions^15^. Although their biological roles remain unclear, one study has suggested that specific tRNA genes are involved in the transcriptional silencing of the amoebapore-A gene^16^. Short tandem repeats between tRNA genes exhibit extensive copy-number polymorphisms, and serve as genetic markers for *E. histolytica* strain typing^17^.

Another distinctive feature of *Entamoeba* genomes is their variable ploidy. Pulsed-field gel electrophoresis of *E. histolytica* has indicated aneuploidy in a broad range of strains^18^, and nuclear DNA content in trophozoites fluctuates markedly under different culture conditions^19^. Genomic analyses have confirmed diverse patterns of aneuploidy across eleven clinical isolates and dynamic changes in ploidy over time^14^. Unlike tumor cells or *Leishmania*, changes in *E. histolytica* ploidy do not correlate with gene expression levels, implying that only a subset of chromosomes may be transcriptionally active^14^.

To further improve the quality of genome assemblies of *Entamoeba* and our understanding of their genomic architectures, we performed *de novo* genome assembly and annotation of the reference strain HM-1:IMSS cl6 of *E. histolytica* and the recently isolated *E. dispar* strain NA442 using PacBio HiFi sequencing. We present their chromosome-level genome assemblies, which provide enhanced insights into *Entamoeba*-specific genomic features, including repetitive structures, tRNA arrays, and aneuploidy.

## Materials and Methods

### Sample collection and cell culture

Trophozoites of *E. histolytica* HM-1:IMSS cl6 2001 were cultured in BI-S-33 medium with Penicillin-Streptomycin-Amphotericin B (Gibco, Grand Island, NY, USA). This strain was originally frozen in 2001 and subsequently refrozen on September 28, 2017. For this study, the 2017 cryopreserved stock was revived on July 9, 2024. The frozen stock was made by the following protocol. One 6-mL tube of *E. histolytica* culture in the semi-logarithmic phase containing around 1×10L trophozoites was placed on ice for 5 minutes, then centrifuged to collect the trophozoites. Add 1 mL of cryopreservation medium [CPM: 5 mL Cell Banker 2 (Zenogen Pharma, Fukushima, Japan), supplemented with 1.14 mg ferric ammonium citrate, 150 mg yeast extract, 5 mg cysteine hydrochloride monohydrate, and 1 mg ascorbic acid] and transfer to two cryotubes (0.5 mL/tube). Put the tubes in a Bicell biofreezing vessel, store at -80°C overnight, then transferred to liquid nitrogen. For reviving, amoebae were rapidly thawed in a 37°C water bath, transfer to 6-mL glass tube with 5 mL of BI-S-33 medium. *E. dispar* NA442 obtained from an outpatient at the AIDS Clinical Center (ACC) of the National Center for Global Health and Medicine (NCGM) in 2023 and maintained in yeast extract-iron-maltose-dihydroxy-acetone-serum (YIMDHA-S) medium^20,21^ supplemented with *Crithidia fasciculata* and *Pseudomonas aeruginosa*. The procedure and methods for isolating and establishing this strain will be explained elsewhere (Saito-Nakano et al., in press).

### Genome sequencing

To isolate genomic DNA, total of 6.7×10^7^ and 3×10^7^ of *E. histolytica* HM-1:IMSS cl6 2001 and *E. dispar* NA442 were prepared. *Entamoeba* trophozoites were washed in PBS then lysed using Cell Lysis Buffer C1 (QIAGEN, Hilden, Germany). The resulting lysate was centrifuged at 1300 × *g* for 10 min at 4°C. The pellet containing nuclei was washed and reacted in ice cold buffer C1 before centrifugation at 1300 × *g* for 10 min at 4°C. The washed pellet was treated with Buffer G2 (QIAGEN), then mixed carefully by inversion followed by treatment with Protease K (QIAGEN) and RNase (QIAGEN). The mixture was incubated for 1 h at 50 °C. Then 1:1 TE-saturated phenol-chloroform was added to the solution then gently mixed by inversion and incubated for 10 min on ice. Afterwards, the mixture was centrifuged at 9,600 rpm, 10 min, 4 °C. The upper phase was transferred on to a new tube and reacted with chloroform, incubated for 2 min on ice, then centrifuged at 9600 rpm for 10 min at 4 °C. The upper phase was collected and treated with Pellet Paint (Novagen, Madison, WI, US), 3M sodium acetate (pH 5.2) and ethanol to precipitate DNA.

The genomic DNA pellet was washed with 70% ethanol. After removal of residual ethanol, the pellet was dissolved in TE buffer before storage at -80 °C. The resultant genomic DNA from HM-1:IMSS (3.5 μg) was directly used for library production. The DNA quality was assessed by electrophoresis using the 5200 Fragment Analyzer System and the Agilent HS Genomic DNA 50 kb Kit (Agilent Technologies, Santa Clara, CA, USA). DNA was purified using DNA Clean Beads (1.8× sample volume; MGI Tech Co., Ltd., Shenzhen, Guangdong, China) and the DNeasy PowerClean Pro Cleanup Kit (QIAGEN). DNA was sheared to an average fragment size of approximately 10–25 kb using the Megaruptor 3 (Diagenode, Liège, Belgium). The fragmented DNA was further concentrated using DNA Clean Beads (1.8× sample volume; MGI Tech Co., Ltd.). Library preparation was performed using the SMRTbell Prep Kit 3.0 (PacBio, Menlo Park, CA, USA) according to the default workflow. For NA442 strain, SMRTbell gDNA Sample Amplification Kit (PacBio) was also used (final 3.6 μg). Polymerase complexes were formed with the prepared libraries using the Revio Polymerase Kit (PacBio) following the manufacturer’s instructions, and sequencing was performed on the Revio system (PacBio).

### Genome assembly of *E. histolytica* HM-1:IMSS cl6

The reads were filtered to a minimum length of 5,000 bp using seqkit (v2.8.2)^22^ and then assembled with hifiasm (v0.19.9-r616)^23^ using the following command: hifiasm -f0 -z20 -l0 - o !{id} -t !{task.cpus} --h1 !{hicr1} --h2 !{hicr2} !{fastq} 2> log_!{id}.txt. For *E. histolytica*, we additionally incorporated publicly available Hi-C reads from the same strain in our previous study (DRR213876) improving assembly accuracy^14^. The raw assembly graph was exported to FASTA and evaluated with QUAST (v5.2.0)^24^, yielding an initial assembly of 311 contigs total of 36,222,116 bp. However, visual inspection using Bandage (v0.9.0)^25^ revealed numerous low-coverage junk contigs. By restricting output to contigs with coverage between 30× and 999,999×, we reduced the assembly to 63 contigs total of 27,581,176 bp. We then remapped the HiFi reads to these 63 contigs and manually removed several redundant contigs identified by low mapping quality scores, resulting in a final set of 35 chromosomal contigs plus one plasmid-derived fragment accounting 26,238,933 bp. The full assembly workflow is publicly available on GitHub (https://github.com/TKSjp/assembly_nf_oss).

### Genome assembly of *E. dispar* NA442

Contrary to *E. histolytica* HM-1:IMSS cl6 2001, *E. dispar* NA442 is maintained in xenic culture with *P. aeruginosa* and *C. fasciculata*. For the initial assembly, we first co-assembled the mixed genome of *P. aeruginosa*, *C. fasciculata*, and *E. dispar*, and then used the non-*E. dispar* assemblies to remove contaminant reads. In this step, metaFlye (v2.9.4-b1799)^26^ produced a more contiguous *P. aeruginosa* assembly (7 contigs total of 6,722,379 bp) than hifiasm. Reads corresponding to *C. fasciculata* were nearly absent, and no contigs were recovered. Finally, we trimmed *E. dispar* reads to a minimum length of 5,000 bp with seqkit and filtered them against the *P. aeruginosa* assembly using minimap2 (v2.28)^27^, then assembled the filtered reads with hifiasm (hifiasm -f0 -z20 -l0 -o !{id} - t !{task.cpus} !{fastq} 2> log_!{id}.txt). QUAST evaluation of this assembly yielded 2,182 contigs totaling 79,327,905 bp. However, as with *E. histolytica*, most contigs were extremely low-coverage junk contigs when visualized in Bandage. The mean coverage was ∼100–150×, so by extracting contigs with coverage between 40× and 999,999×, we reduced the set to 52 contigs total of 27,801,265 bp. After manually removing a few remaining redundant contigs identified by low mapping quality score, we obtained 35 chromosomal contigs plus one plasmid-derived fragment, with a total length of 27,015,509 bp.

### Genome annotation of *E. histolytica* and *E. dispar*

Prior to annotation, repeats were masked by running RepeatModeler^28^ and RepeatMasker^29^ (RepeatModeler -database ${id} -threads ${task.cpus} -LTRStruct followed by RepeatMasker -pa 2 -dir ${id}_repeatmasker -lib RM/consensus_plus.fa ${fna}) using the consensus sequences from RepeatModeler and all currently available *Entamoeba* LINE, SINE, mariner, ERE1, and ERE2 elements^13^. The soft-masked FASTA was then subjected to tRNA gene prediction with tRNAscan-SE (v.2.0.12; tRNAscan-SE --thread ${task.cpus} - ${mode} --bed ${id}_${mode}.bed --fasta ${id}_${mode}.fasta --stats ${id}_${mode}.stats ${fna})^30^. We used “O” mode for maximal sensitivity and “E” mode to distinguish initiation methionine (iMet) versus elongation methionine (eMet). Other ncRNAs were identified with Infernal (1.1.5) via cmscan --cut_ga --tblout ${id}.tblout --fmt 2 --cpu ${task.cpus} -o ${id}.out ${params.infernal_cm} ${fna}^31^. Coding genes were predicted by running BRAKER3 (braker.pl --genome=${fna} --prot_seq=${params.braker_protseqfasta} --workingdir=intermediate --threads ${task.cpus} --gff3 --species ${params.braker_species} --useexisting --AUGUSTUS_CONFIG_PATH ${params.augustus_config_path}) with *E. histolytica* proteome in AmoebaDB release 63 for the --prot_seq^32,33^. Finally, predicted ORFs were annotated by DIAMOND (2.1.12; diamond blastp -d ${params.diamond_db} -q ${faa} -o ${id}.tsv) against the UniRef90 database^34,35^. The following options were used: --threads ${task.cpus} --sensitive --evalue 1e-5 --max-target-seqs 10 --query-cover 50 --subject-cover 50 --outfmt 6 qseqid sseqid pident length mismatch gapopen qstart qend sstart send evalue bitscore stitle. The complete workflow is available on GitHub (https://github.com/TKSjp/annotation_nf_oss).

### Genome comparison

For chromosome-scale synteny visualization, we used Circos (v0.69-9)^36^. First, we created a BLAST database for *E. dispar* with BLAST+ (2.16.0+; makeblastdb -dbtype nucl -in NA442.fna -out NA442.fna -hash_index)^37^. We then performed BLASTn mapping (blastn - query cl6.fna -db NA442.fna -outfmt “6 qseqid qstart qend sseqid sstart send pident length” -num_threads 10 -evalue 1e-10 -max_target_seqs 30 > AB.blast6) to map the *E. histolytica* contigs to the *E. dispar* contigs. Alignments were converted and filtered with an AWK script, retaining hits ≥2,000 bp and ≥90% identity (awk ‘$8 ≥ 2000 && $7 ≥ 90 { print “cl6_“$1“\t“$2-1“\t“$3“\tNA442_“$4“\t“$5-1“\t“$6}’ AB.blast6 > links_original.txt). Amino acid-level synteny was analyzed with MCScanX (v1.0.0)^38^. GFF files from both *E. histolytica* and *E. dispar* were concatenated, after which a protein BLAST database was constructed and the all-versus-all search was run (blastp -query cat.faa -db cat.faa -evalue 1e-50 - qcov_hsp_perc 90 -max_target_seqs 5 -outfmt 6 -num_threads 16). Results were visualized in the SynVisio web application^39^ by selecting chromosomes 1 through 15 in the filter panel. Finally, proteome overlap between *E. histolytica* and *E. dispar* was assessed using the OrthoVenn3 web server^40^, with the default OrthoMCL algorithm, an E-value cutoff of 1e-2, and an inflation value of 1.5.

### Ideogram

We used RIdeogram (0.2.2)^41^ to draw ideograms. First, we generated a karyotype.tsv file from the assembled genome sequence. We then used the mRNA features in the GFF file to calculate and plot gene density tracks. Finally, we extracted the tRNA, 5S rRNA, and Metazoa SRP (7SL RNA) annotations from the GFF and overlaid these features onto the ideogram using RIdeogram. Additionally, regions with excessively dense annotations like tRNA genes were thinned using a custom Python script, which identifies clusters of features spaced less than 10 kb apart and retains only a representative subset to avoid overplotting.

### Ploidy comparison

The HiFi reads trimmed with fastplong (0.2.2)^42^ were mapped to the genome assembly using minimap2, and the resulting alignments were converted into sorted BAM format with Samtools (1.20)^43^. Read depth was then calculated using bamCoverage implemented in DeepTools^44^, retaining only alignments with a mapping quality greater than 50 (bamCoverage --bam input.bam -o output.bed -of bedgraph --binSize=10000 --minMappingQuality 50 --numberOfProcessors=max). The resulting BED file was imported into R, and read depth profiles were visualized using ggplot2.

## Results

### Comparison of genome structures between *E. histolytica* HM-1:IMSS cl6 and *E. dispar* NA442

Using PacBio Revio HiFi sequencing, we performed de-novo, high-quality genome assemblies for *E. histolytica* and *E. dispar* (Fig. 1A). Sequencing produced 7.5 Gb of reads for *E. histolytica* and 6.8 Gb for *E. dispar*, which decreased, after trimming, to 7.2 Gb and 6.6 Gb, respectively. For *E. histolytica* we supplemented the HiFi reads with previously published Hi-C data from the same strain^14^ and assembled both genomes with hifiasm. Low-depth contigs (<30×) were discarded in Bandage, and redundant contigs were manually removed after read-mapping inspection. The final assemblies of the *E. histolytica* genome comprise 36 chromosome-size contigs plus a single plasmid totaling 26 Mb, while those for the *E. dispar* genome comprise 35 chromosome-size contigs plus one plasmid totaling 27 Mb for. Repeat masking with RepeatModeler and RepeatMasker revealed that 37.6% and 38.1% of the *E. histolytica* and *E. dispar* genome, respectively, consist of repetitive elements. Annotation using tRNAscan-SE, Infernal, BRAKER3, and DIAMOND (UniRef90) identified 1,474 tRNA genes, 134 ncRNA genes, and 10,008 coding genes in *E. histolytica*, and 1,462 tRNA genes, 167 ncRNA genes, and 9,803 coding genes in *E. dispar*. Actual gene number for tRNA are likely higher, as array regions are not fully resolved. In addition to the known intron-bearing tRNA^Tyr^(GUA) and tRNA^Ile^(UAU)^45^, we discovered additional intron-bearing tRNA genes including tRNA^Val^(UAC), tRNA^Ser^(ACU), tRNA^Leu^(UAA), and tRNA^Tyr^(AUA).

**Figure 1.**
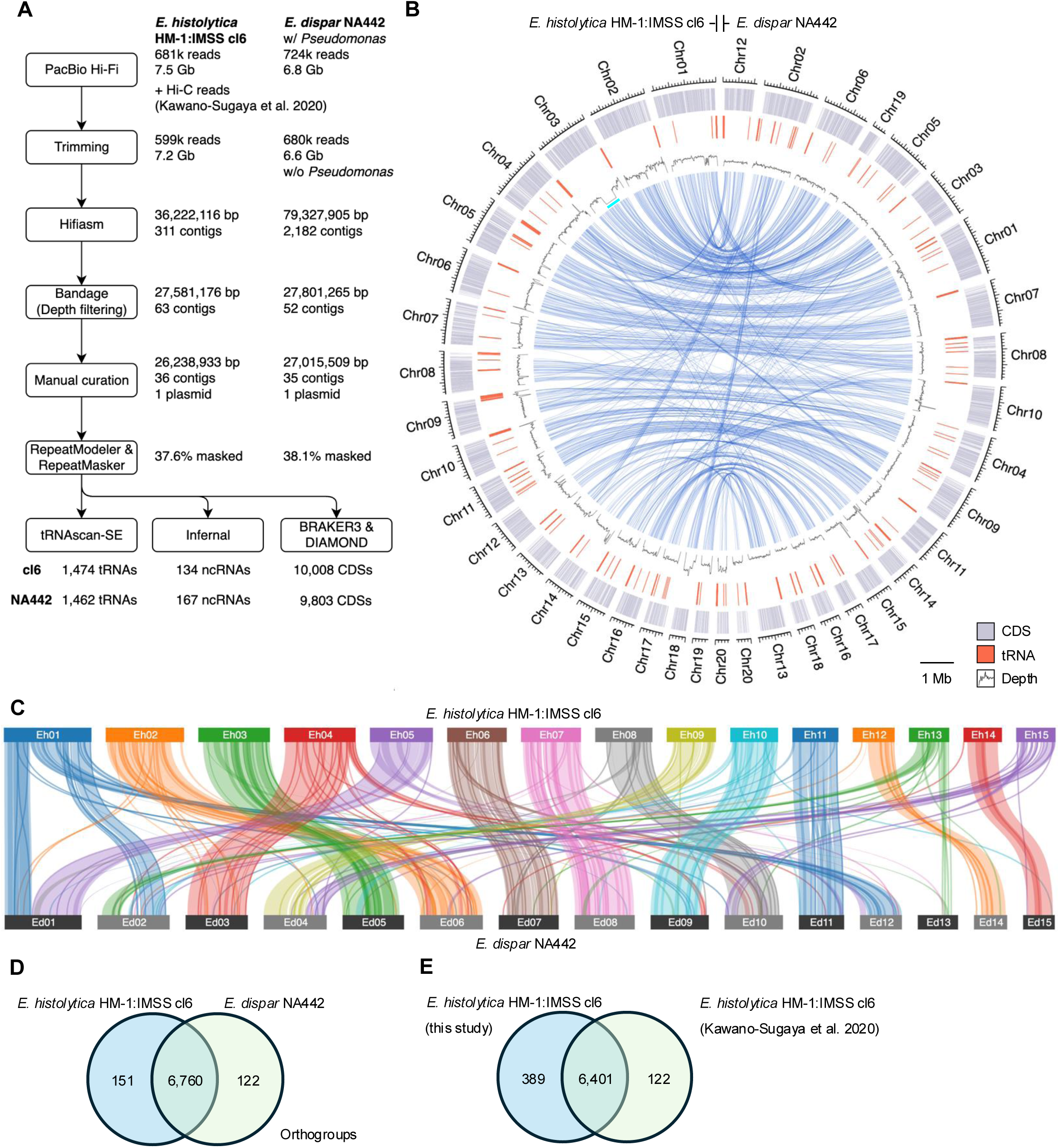
Genome analysis and structural comparison. (**A**) Workflow and summary statistics of the genome analysis. (**B**) Chromosome organizations representing both genomes of *E. histolytica* HM-1:IMSS cl6 2001 and *E. dispar* NA442. Read depth was calculated by mapping PacBio HiFi reads with minimap2 (MAPQ>50). Relationships among contigs were aligned by BLASTn. For clarity, chromosomes 1–20 are shown. Cyan region marks a long repetitive segment lacking uniquely mapped reads (Fig. S1). (**C**) Amino–acid–level synteny identified by MCScanX and visualized by SynVisio. For clarity, chromosomes 1–15 are shown. (**D**) Venn diagram showing the number of shared and species-specific orthogroups between *E. histolytica* HM-1:IMSS cl6 2001 and *E. dispar* NA442. (**E**) Venn diagram showing the number of shared and assembly-specific orthogroups between *E. histolytica* HM-1:IMSS cl6 2001 in this study and Kawano-Sugaya et al. 2020^14^.

A whole-genome comparison generated with Circos (Fig. 1B) revealed a strong chromosome-scale similarity. Amino-acid sequence-level synteny visualized with MCScanX and SynVisio confirmed that most chromosomes retain clear collinearity (Fig. 1C; e.g., Eh02–Eh04, Eh06–Eh08, Eh10–Eh11, Eh14 and their counterparts Ed03, Ed05–Ed11, Ed15). In several cases, single *E. histolytica* chromosomes correspond to two separate *E. dispar* chromosomes (e.g., Eh01 with Ed02 and Ed12) or vice versa, indicating block-level structural rearrangements. Despite these rearrangements, overall gene content remained highly conserved: OrthoVenn3 identified 6,760 shared orthogroups, with only 151 Eh-specific and 122 Ed-specific groups (Fig. 1D). Thus, both nucleotide sequences and encoded proteins show extensive conservation between *E. histolytica* and *E. dispar*.

We observed discrepancies when comparing our annotations to previous reports. These discrepancies are most likely attributable to different gene prediction and annotation methods used. Our current gene counts for *E. histolytica* (10,008) and for *E. dispar* (9,803) exceed earlier published numbers by 992–1,857 genes (*E. histolytica*: 8,151^8^ and 8,734^14^; *E. dispar*: 8,811^9^). An OrthoVenn3 comparison with the previous annotation^14^ identified 389 orthogroups unique to our assembly (Fig. 1E), which is greater than species-level differences observed between *E. histolytica* versus *E. dispar* (151; Fig. 1D). Former work^14^ used Companion with AUGUSTUS^46^ for gene prediction, whereas we used BRAKER3, suggesting that differences in annotation pipelines are a major contributor to our higher gene counts. Incomplete repeat masking also seems to contribute gene counts: 34 non-LTR retrotransposon reverse-transcriptase genes in *E. histolytica* and 18 in *E. dispar* remained unmasked, and additional unmasked transposon-derived ORFs may have inflated our total genes. Thus, advances in annotation tools likely contributed to the increased gene numbers, and further improvements in preprocessing and repeat-handling protocols in the gene prediction and annotation pipeline are essential for ensuring consistency with past annotations.

### tRNA array locations among chromosomes

While overall chromosome structures were conserved between *E. histolytica* and *E. dispar* (Fig. 1), we next asked whether the polymorphism and distribution of tRNA arrays also show conservation between the two species. Previous analyses of *Entamoeba* tRNA arrays were based on SangerLsequenced genomes^8,9^. These previous studies identified 25 distinct tRNA arrays in *E. histolytica* and 24 in *E. dispar* (all but the NK1 array containing Asn^GTT and Lys^CTT), with conserved array organizations between the two species^9^.

We visualized the chromosomal locations of all tRNA arrays, identified based on our longLread assemblies and tRNAscan-SE^41^ (Fig. 2). We identified 28 distinct tRNA arrays in both *E. histolytica* and *E. dispar* (Fig. 2A, B). Some arrays include 5S rRNA or snRNA, which is consistent with earlier reports^9^. Although NK1 and NK2 had been reported only in *E. histolytica*, and NK1 was predicted to be absent in *E. dispar*^9^, we detected two of NK arrays on separate chromosomes in both species (Fig. 2B, blue). Moreover, we discovered that the YE (Tyr^GTA, Glu^TTC), WI (Trp^CCA, Ile^AAT), and VF (Val^GAC, Phe^GAA) arrays each are present at two locations; we have denoted these pairs as YE1/YE2, WI1/WI2, and VF1/VF2 in order of decreasing chromosome size (Fig. 2A, magenta).

**Figure 2.**
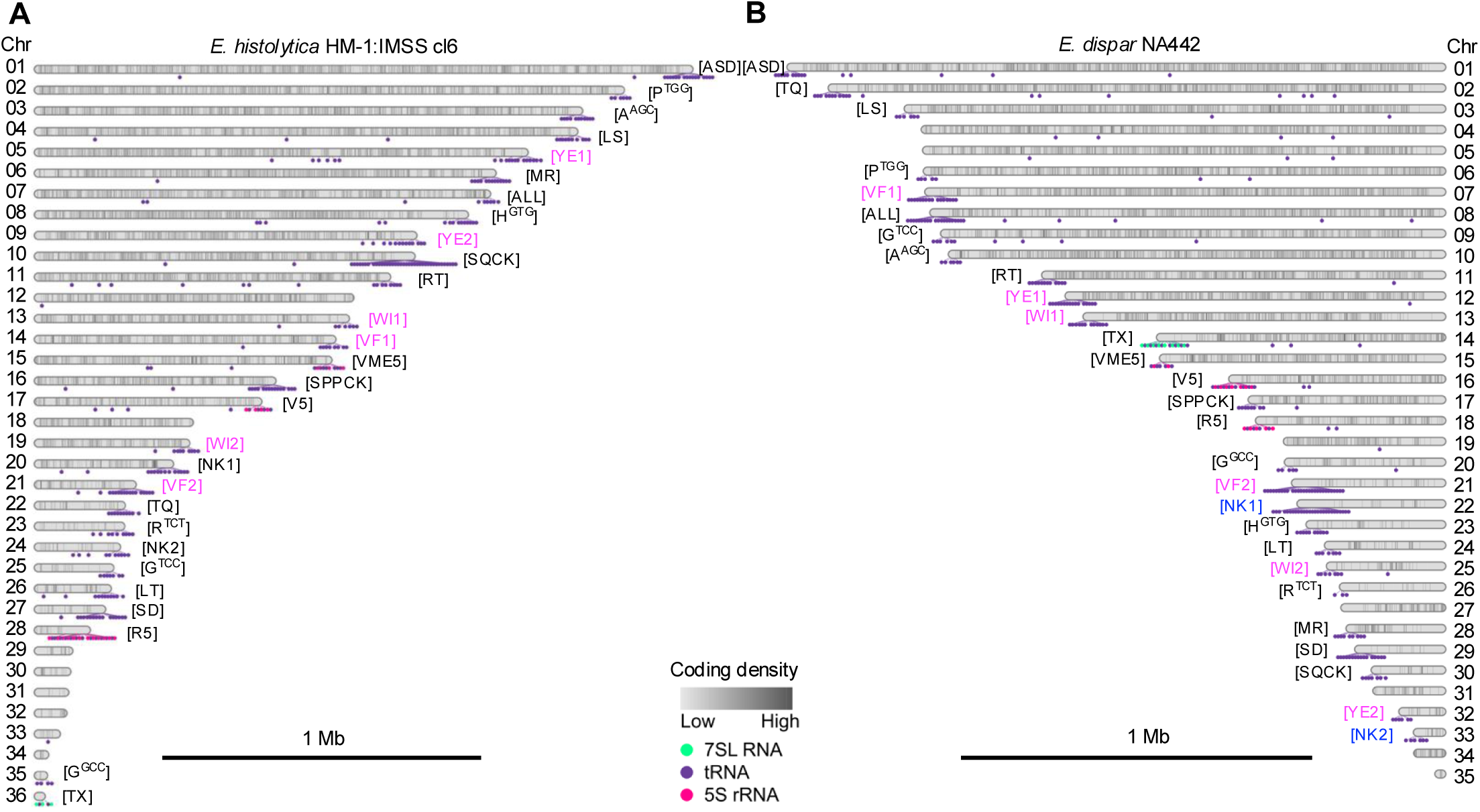
Ideogram and tRNA array. (**A**) Ideogram of *E. histolytica* HM-1:IMSS cl6 with the positions of all tRNA arrays. Cyan dots denote 7SL RNA, purple circles denote tRNAs, and magenta circles denote 5S rRNA sequences. To avoid overplotting, marker density in highly clustered regions was reduced to 20% of the true number. Each array is labelled with its canonical tRNA-array name; arrays newly identified in this study are written in magenta. (**B**) Ideogram of *E. dispar* NA442 and its tRNA arrays. The NK arrays, one of which has been identified in this study, are written in blue.

Overall, the types of tRNA arrays in *E. dispar* matches those in *E. histolytica*. The same tRNA arrays are present on homologous chromosomes between the two *Entamoeba* species (e.g. Eh01–Ed01; Eh02–Ed06; Eh04–Ed03; Eh07–Ed08; Eh11–Ed11). However, exceptions were also observed: a paired homologous chromosomes (Eh03–Ed05; Eh06–Ed07) contain different tRNA arrays, suggesting that certain tRNA arrays have undergone inter-chromosomal translocation.

### Non-LTR retrotransposons in *Entamoeba*

We found that over 35% of both the *E. histolytica* and *E. dispar* genomes consist of repetitive elements, reflecting massive expansion of transposable elements in *Entamoeba*. Although SINEs are only a few hundred base pairs long, the three *E. histolytica* LINE families are relatively large^47^ (EhLINE1: 4,804 bp, EhLINE2: 4,722 bp, and EhLINE3: 4,845 bp), which likely contributes to the difficulty of resolving these repeats in earlier Sanger-based assemblies. To investigate these repeat elements in depth, we used RepeatModeler and RepeatMasker using known *Entamoeba* LINE, SINE, ERE1, and ERE2 sequences as references to profile transposons in *E. histolytica* and *E. dispar*. However, RepeatMasker called none of the known SINEs, LINEs, LTRs, or DNA elements detectable with default parameter settings in both species. In *E. histolytica*, 8,509 unclassified elements (34.9% of the genome), 9,771 simple repeats (2.0%), and 3,334 low-complexity regions (0.7%) were masked. Similarly, in *E. dispar*, 8,597 unclassified elements (35.0%), 10,915 simple repeats (2.2%), and 3,729 low-complexity regions (0.8%) were masked. The unclassified category includes previously described *Entamoeba* LINEs, SINEs, ERE1s, and ERE2s. From these unclassified elements, we recovered 113 LINEs, 306 SINEs, 1 mariner element, 430 ERE1s, and 416 ERE2s (1,266 total) in *E. histolytica*, whereas *E. dispar* contained 76 LINEs, 80 SINEs, 171 ERE1s, and 10 ERE2s (337 total)—only 26.6% of the number detected in *E. histolytica*. Finally, we identified an additional 112 and 96 previously non-discovered repeat families in *E. histolytica* and *E. dispar*, respectively, which together account for the majority of the remaining repetitive content.

### Different ploidy patterns between *E. histolytica* and *E. dispar*

It was previously shown that eleven *E. histolytica* clinical isolates exhibited distinct chromosome- or segment-dependent ploidy patterns, with some regions reaching as high as septaploid^14^. However, those inferences were based on short-read mapping data. To overcome such ambiguity inherited by error-prone sequencing methods, we aimed to visualize more accurate ploidy patterns by mapping our high-quality HiFi reads back to the new assemblies. Across chromosomes 1–20, uniquely mapped HiFi reads yielded an average depth of approximately 500× in *E. histolytica* (Fig. 3A). Notably, four regions on Chr2, Chr10, Chr18, and Chr20 reached depths of ∼600× to 750× or higher; if a baseline of 500× represents tetraploidy, these elevated depths imply local copy numbers of five, six, or higher numbers. In addition, Chr2 and Chr10 show markedly different average depths between their first and second halves, suggesting intrachromosomal variation in ploidy. Conversely, a region highlighted by a blue arrow displays an average depth of ∼250×, roughly half the genome-wide mean, supporting the assumption that this region is diploid. These observations are largely in agreement with our previous findings^14^. Note that cyan-marked segments on Chr3 correspond to large repetitive regions that lack uniquely mappable reads (Fig. S1).

**Figure 3.**
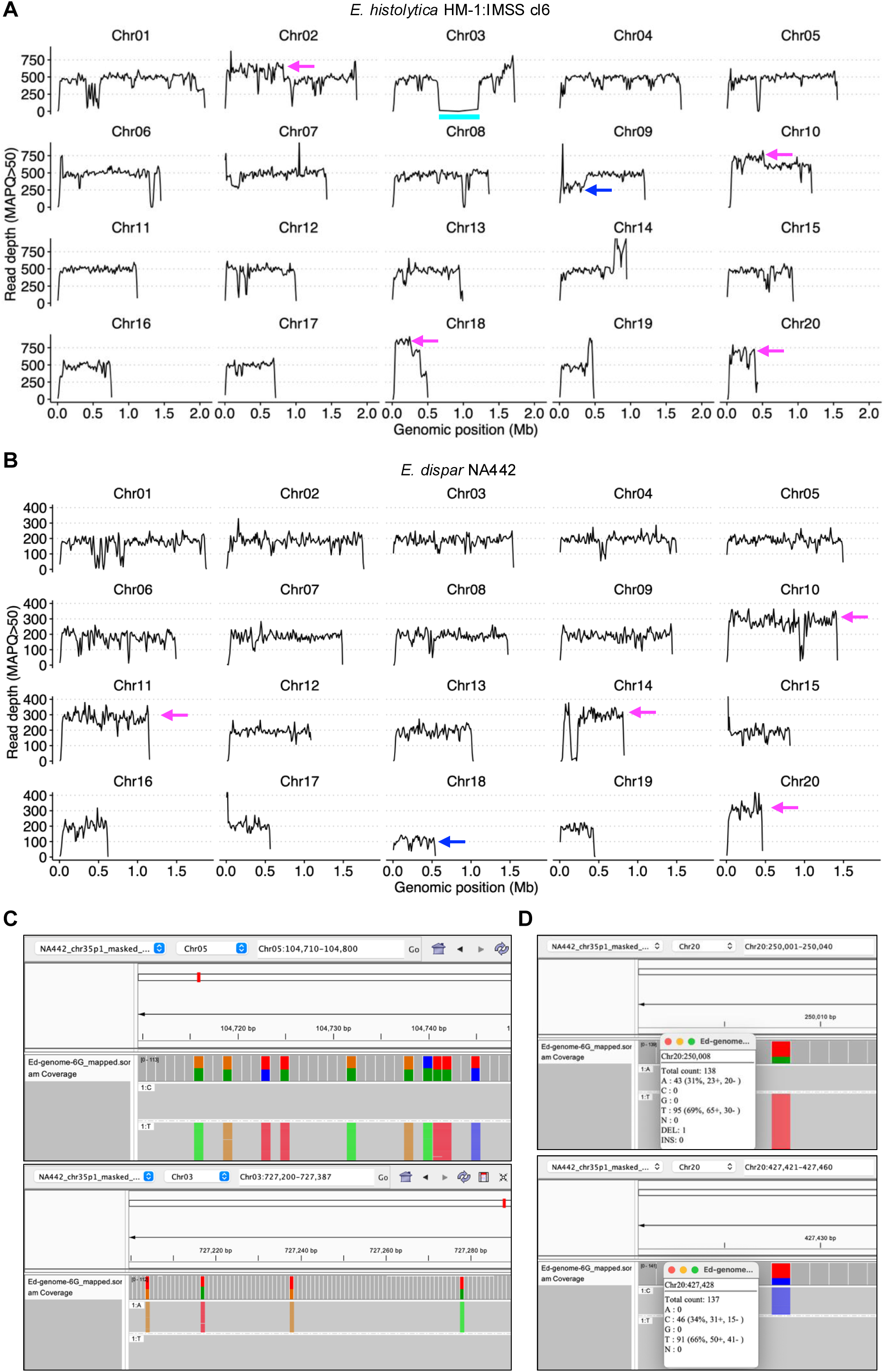
Ploidy variation in *E. histolytica* HM-1:IMSS cl6 and *E. dispar* NA442. (**A**) Read-depth profiles and inferred ploidy for each chromosome of *E. histolytica*. Magenta regions indicate depth above the genome-wide average; blue regions indicate depth below the average; cyan region mark a long repetitive segment lacking uniquely mapped reads (Fig. S1). (**B**) Read-depth profiles and inferred ploidy for each chromosome of *E. dispar*. Chromosome 18 exhibits an average depth of approximately ∼100×, consistent with monoploidy. (**C**) Example of heterozygosity in *E. dispar* showing allele frequencies are 1:1, consistent with a diploid region. (**D**) Example of heterozygosity in *E. dispar* showing allele frequencies are 1:2, consistent with a triploid region.

In contrast, in *E. dispar*, the average unique-read depth was approximately 200× (Fig. 3B). Four genomic regions showed increased depths of ∼300× (magenta), whereas one region exhibited a markedly lower depth of ∼100× (blue). Unlike *E. histolytica*, none of the *E. dispar* chromosomes displayed intra-chromosomal variations in read-coverage depth between segments on each chromosome. Although no previous experimental evidence, such as pulsed-field gel electrophoresis plus Southern blot analysis, DNA melting assay, and flow cytometry DNA content assay, to estimate ploidy are available for *E. dispar,* the observed depth distribution strongly infers a predominant diploidy, with a few segments of triploidy and a single region of monoploidy (Fig. 3B). Further support for diploidy in *E. dispar* comes from heterozygosity patterns. In *E. histolytica*, heterozygous sites occur genome-wide, showing allele-frequency peaks at 25:75 and 50:50, consistent with a predominant tetraploid state¹³. In the *E. dispar* genome, most heterozygous sites clearly showed the allele-frequency ratios close to 50:50 (Fig. 3C). Two exceptions were found on Chr20, which was inferred to be triploid based on read depth; these sites displayed allele-frequency ratios of 31:69 and 34:66 (Fig. 3D). Together, these data indicate that *E. dispar* possesses a largely homozygous, diploid genome, with Chr20 representing a notable triploid exception.

## Discussion

The *E. histolytica* genome is highly repetitive, and even with long-read sequencing it has been challenging to resolve the numerous repeat sequences and build a high-quality genome assembly. In our previous work, we scaffolded the genome into 38 but still separated into 69 contigs^14^. To achieve further improvement, it was essential to assemble the genome without scaffolding and to pinpoint the exact locations of the tRNA arrays—an effort that required reconstructing each chromosome as a single contiguous sequence. In this study, we produced assemblies comprising 36 single-contig chromosomes for *E. histolytica* and 35 for *E. dispar*. Among these, eight contigs in *E. histolytica* and fourteen contigs in *E. dispar* were shorter than 300 kb, which is notably smaller than the chromosome sizes estimated by pulsed-field gel electrophoresis (PFGE) in Willhoeft et al. (1999), who reported 31–35 chromosomes in *E. histolytica* ranging in size from 0.3 to 2.2 Mb^18^. This apparent discrepancy may reflect the lower size detection limit of their PFGE system, whose smallest resolvable band was approximately 225 kb, substantially larger than the smallest chromosomes we identified. Their rotating-field electrophoresis setup also suggested the presence of chromosomes smaller than 225 kb. Additionally, natural karyotypic variation among *E. histolytica* isolates may further contribute to differences between their PFGE-based estimates and our estimates by long-read assemblies^18^.

We visualized chromosome-scale alignments using Circos and found that chromosome organization is largely conserved between *E. histolytica* and *E. dispar*. Given the minimal differences in overall gene content, the phenotypic divergence between the two species is likely driven not by large-scale genome rearrangements but rather by gene-level alterations, such as changes in gene regulation and the accumulation of small nucleotide substitutions that lead to functional or expression differences.

Several new findings on tRNA array presented by this study are largely different from what was previously described. Notably, we detected two NK arrays in *E. dispar*, whereas only one had been described previously. Likewise, the YE, WI, and VF arrays, which have been thought to exist as a single locus, were found as two arrays in both species. These observations may reflect divergence of our laboratory strain, *E. histolytica* HM-1:IMSS cl6, from the original HM-1:IMSS or may capture strain-level variation between our *E. dispar* NA442 and the previously sequenced SAW760. Alternatively, improvements in long-read sequencing and annotation methods may have enabled discoveries of common genetic features that were overlooked in earlier studies.

We also took advantage of alternative tRNAscan-SE models (the “Other Organellar” mode in addition to Eukaryote mode) to identify intron-containing tRNAs that had not been reported before. We discovered additional intron-bearing tRNA genes including tRNA^Val^(UAC), tRNA^Ser^(ACU), tRNA^Leu^(UAA), and tRNA^Tyr^(AUA). Although tRNA^Ile^(UAU) intron splicing in *E. histolytica* is mediated by nuclear RtcB1, some of these newly identified intron-containing tRNAs may instead be processed by the cytosolic RtcB2. Future studies of their biogenesis and function may provide insights into processing pathways and potentially reveal new targets for anti-amoebic drug development.

Despite the high overall nucleotide similarity between the two genomes, their non-LTR retrotransposon repertoires were different. In the present survey, the numbers of LINEs, SINEs, ERE1s and ERE2s identified from *E. dispar* represent only 26.6% of those detected in *E. histolytica*, with SINEs and ERE1s particularly under-represented. Even the counts obtained for *E. histolytica* are lower than those reported in the original profiling of its LINE and SINE families^47^. Whether this discrepancy reflects redundancy in the earlier Sanger-based assemblies or incomplete sensitivity of current repeat-detection pipelines remains unclear. In addition, ten elements classified as ERE2 were identified from *E. dispar*, although ERE2 has previously been considered specific to *E. histolytica*. These copies may have been mis-assigned or could instead represent highly divergent ERE1 elements, a possibility that warrants further investigation.

This work provides the first evidence that *E. dispar* is predominantly diploid, a feature that may make it a more attractive for genome editing than the tetraploid *E. histolytica*. A diploid background halves the number of alleles that must be edited, and the apparent monoploidy of chromosome 18 could further enhance editing efficiency by reducing or preventing homologous-recombination-based repair.

In summary, we presented high-quality, single-contig assemblies for *E. histolytica* and *E. dispar*. Comparative analyses clarified chromosome-scale organization and local structural rearrangements at both nucleotide and protein levels, and revealed previously unrecognized tRNA arrays shared between the species. Mapping of HiFi reads provided refined ploidy estimates, highlighting extensive, chromosome-specific aneuploid variation in *E. histolytica* and a predominantly diploid state in *E. dispar*. These data furnish a robust genomic resource and a framework for future studies in *Entamoeba* genomics.

## Acknowledgements

We thank Koji Watanabe (Tokai University) for support in the initial cultivation and maintenance of the *E. dispar* NA442 strain. We also express our appreciation to Kumiko Shibata (The University of Tokyo) for technical support with *Entamoeba* cultivation and genomic DNA preparation. The super-computing resource was provided by the Human Genome Center, the Institute of Medical Science, and the University of Tokyo.

## Author contributions

Kumiko Nakada-Tsukui (Conceptualization, Funding acquisition, Project administration, Supervision, Writing—review & editing), Tomoyoshi Nozaki (Funding acquisition, Project administration, Writing—review & editing), Tetsuro Kawano-Sugaya (Data curation, Investigation, Methodology, Software, Visualization, Writing—original draft), Seiki Kobayashi (Resources), Akira Kawashima (Resources), Yumiko Saito-Nakano (Resources), and Shinji Izumiyama (Data curation).

## Ethics declarations

The study was conducted in accordance with the Declaration of Helsinki and approved by the Ethics Committee of the National Center for Global Health and Medicine (NCGM-S-004658-02). Written informed consent was obtained from the participant.

## Data availability

All sequence data produced in this study were deposited in the Sequence Read Archive (SRA) and GenBank at the National Center for Biotechnology Information (NCBI). The identifiers in the BioProject and BioSample are PRJNA1365478, PRJNA1365473, SAMN53286930 and SAMN53286573, respectively. The final genome assembly and annotations were deposited in GenBank under accession number JBXEOZ000000000 and JBXEPA000000000.

