## Supplementary figures and images for "Chromosome organization of Entamoeba histolytica and Entamoeba dispar"

### Fig. S1

#### Slide 1
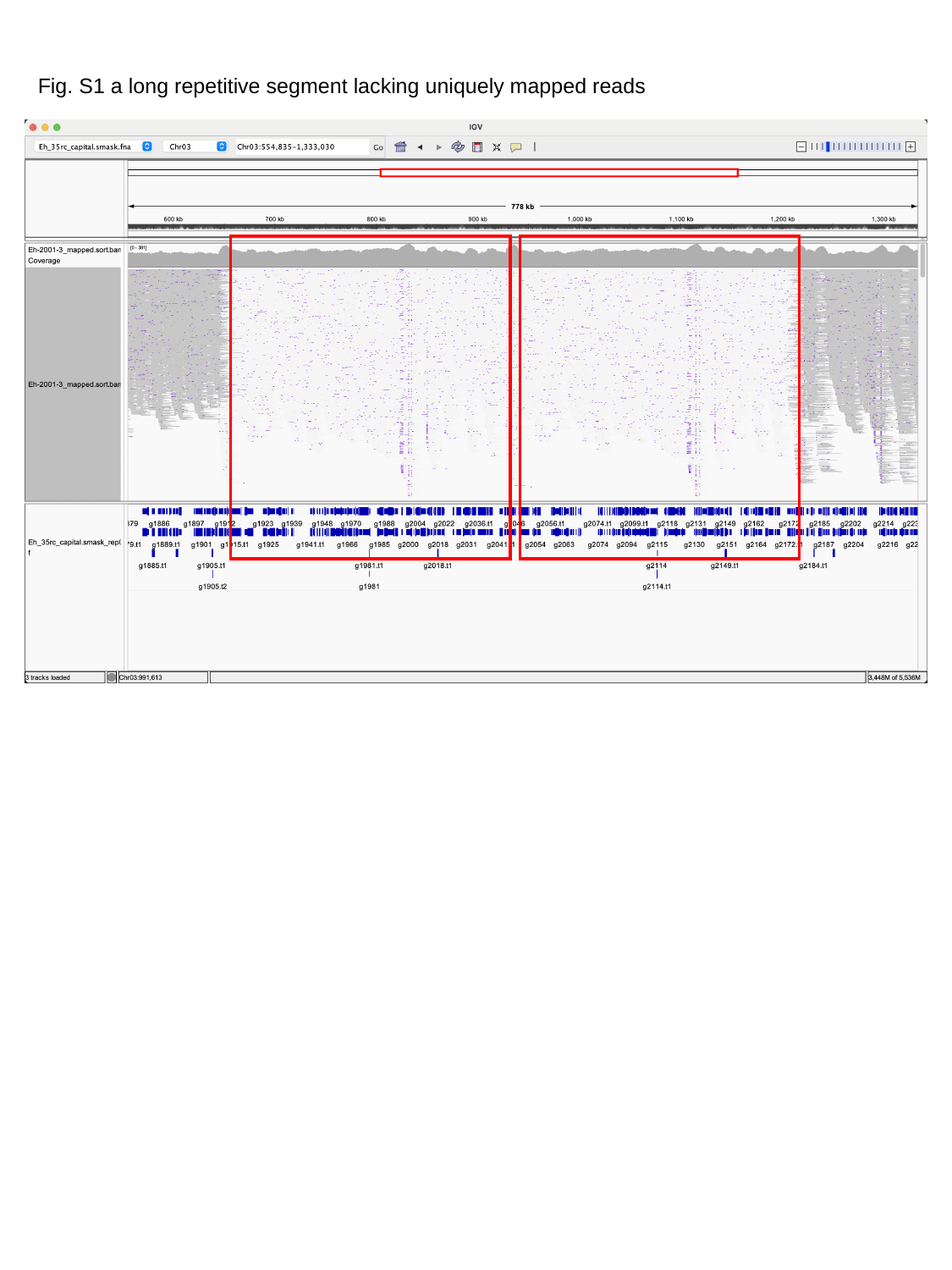

Fig. S1 a long repetitive segment lacking uniquely mapped reads
